# Evolution of the Mineralocorticoid Receptor: Sequence, Structure and Function

**DOI:** 10.1101/098921

**Authors:** Michael E. Baker, Yoshinao Katsu

## Abstract

The mineralocorticoid receptor (MR) is descended from a corticoid receptor (CR), which has descendants in lamprey and hagfish, cyclostomes (jawless fish), a taxon that evolved at the base of the vertebrate line. A distinct MR and GR first appear in cartilaginous fishes (Chondrichthyes), such as sharks, skates, rays and chimaeras. Skate MR has a strong response to corticosteroids that are mineralocorticoids and glucocorticoids in humans. The half-maximal responses (EC50s) for skate MR for the mineralocorticoids aldosterone and 11-deoxycorticosterone are 0.07 nM and 0.03 nM, respectively. EC50s for the glucocorticoids cortisol and corticosterone are 1 nM and 0.09 nM, respectively. The physiological mineralocorticoid in ray-finned fish, which do not synthesize aldosterone, is not fully understood because several 3-ketosteroids, including cortisol, 11-deoxycortisol, corticosterone, 11-deoxycorticosterone and progesterone are transcriptional activators of fish MR. Divergence of the MR and GR in terrestrial vertebrates, which synthesize aldosterone, led to increased selectivity of the MR for aldosterone, coupled with a diminished response to cortisol and corticosterone. Here, we combine sequence analysis of the CR and vertebrate MRs and GRs, analysis of crystal structures of human MR and GR and data on transcriptional activation by 3-ketosteroids of wild-type and mutant MRs and GRs to investigate the evolution of selectivity for 3-ketosteroids by the MR in terrestrial vertebrates and ray-finned fish, as well as the basis for binding of some glucocorticoids by human MR and other vertebrate MRs.

## Introduction

In this special issue of JOE, we celebrate the thirtieth anniversary of cloning of the mineralocorticoid receptor (MR) in the Evans laboratory at the Salk Institute (Arriza, et al. 1987). This was an impressive achievement. Indeed, it was not an easy task, as the MR was the last cloned receptor from the adrenal and sex steroid receptor family, which also includes the glucocorticoid receptor (GR), progesterone receptor (PR), androgen receptor (AR) and estrogen receptor (ER) (Baker, et al. 2015; Evans 1988; Markov, et al. 2009,Baker, 2011 #31). The MR and other steroid receptors belong to the nuclear receptor family, a diverse group of transcription factors that arose in multicellular animals, which have key roles in the physiology of humans and other vertebrates (Baker, et al. 2013; Bridgham, et al. 2010; Huang, et al. 2010; Markov et al. 2009).

The availability of recombinant human MR facilitated the cloning of MRs from a wide variety of vertebrates and an analysis of the evolution of the MR. These MR sequences and those of the GR confirmed the original observation of kinship between the MR and GR (Arriza et al. 1987; Baker, et al. 2007; Bridgham, et al. 2006; Evans 1988). The MR and GR are descended from the corticoid receptor (CR) (Baker et al. 2007; Baker et al. 2015; Bridgham et al. 2006; Thornton 2001). Descendants of the ancestral CR are found in lampreys and hagfish, which are cyclostomes (jawless fish), a taxon that evolved at the base of the vertebrate line (Figure 1) (Osorio and Retaux 2008; Sauka-Spengler and Bronner-Fraser 2008). A distinct MR first appears in cartilaginous fishes (Chondrichthyes), such as sharks, skates, rays and chimaeras (Baker et al. 2015; Bridgham et al. 2006; Carroll, et al. 2008).

**Figure 1.**
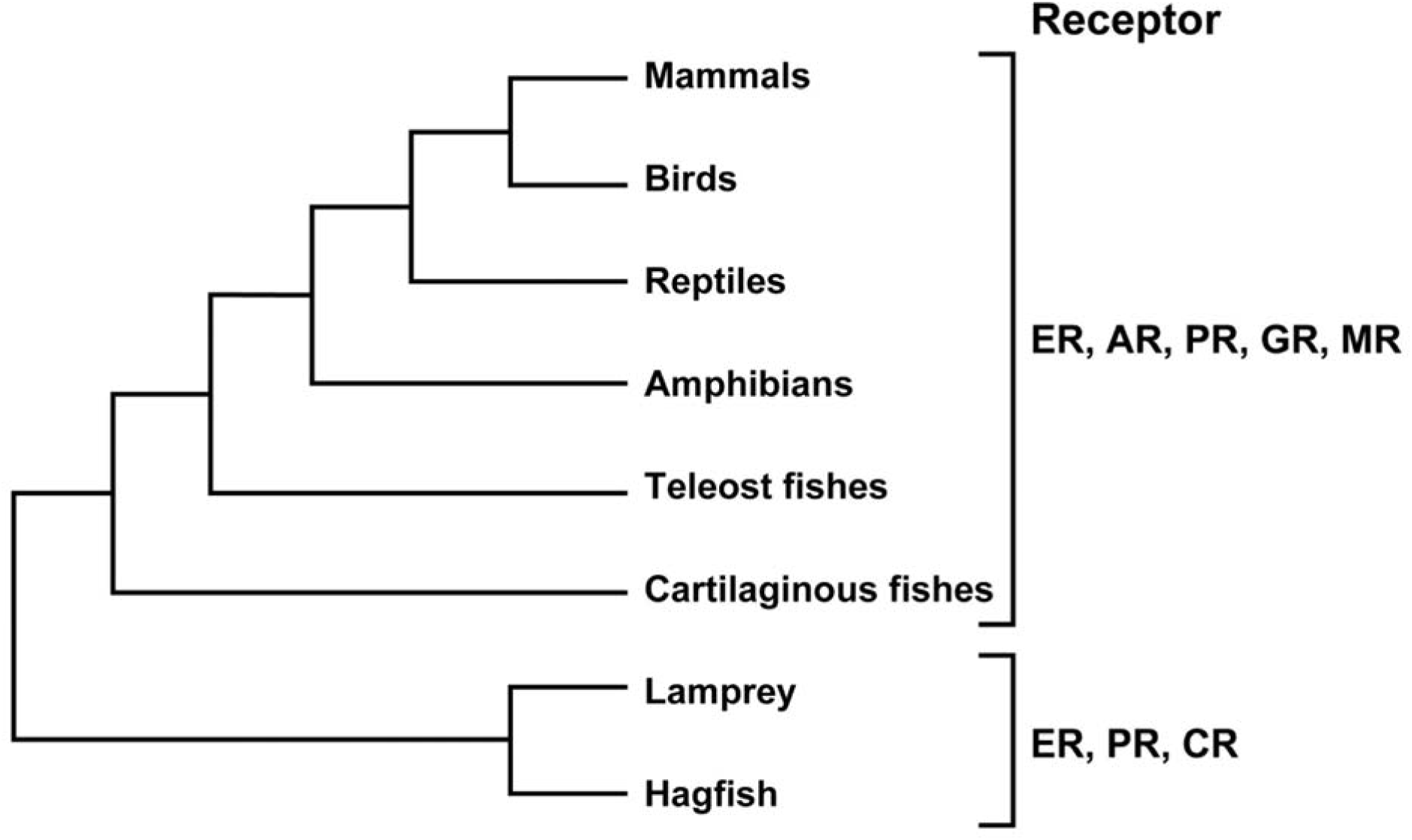
MR, GR, PR, AR and ER in vertebrates. The ER, CR and PR are found in lampreys and hagfish, jawless fishes (cyclostomes) that evolved at the base of the vertebrate line. A separate MR and GR are found in skates, rays, sharks and chimaeras, cartilaginous fishes that are basal jawed vertebrates. The AR first appears in cartilaginous fishes. Land vertebrates are descended from lobe-finned fishes [coelacanths and lungfishes].

Studies with recombinant human MR yielded some unexpected findings. First, Arriza et al. (Arriza et al. 1987) showed that aldosterone (Aldo), cortisol (F), 11-deoxycortisol (S), corticosterone (B), 11-deoxycorticosterone (DOC) and progesterone (Prog) (Figure 2) had similar equilibrium binding constants (Kds) for human MR. This indicated that the steroid binding site on the human MR was not selective for physiological mineralocorticoids (Aldo, DOC) over glucocorticoids (F, B, S) and Prog.

**Figure 2.**
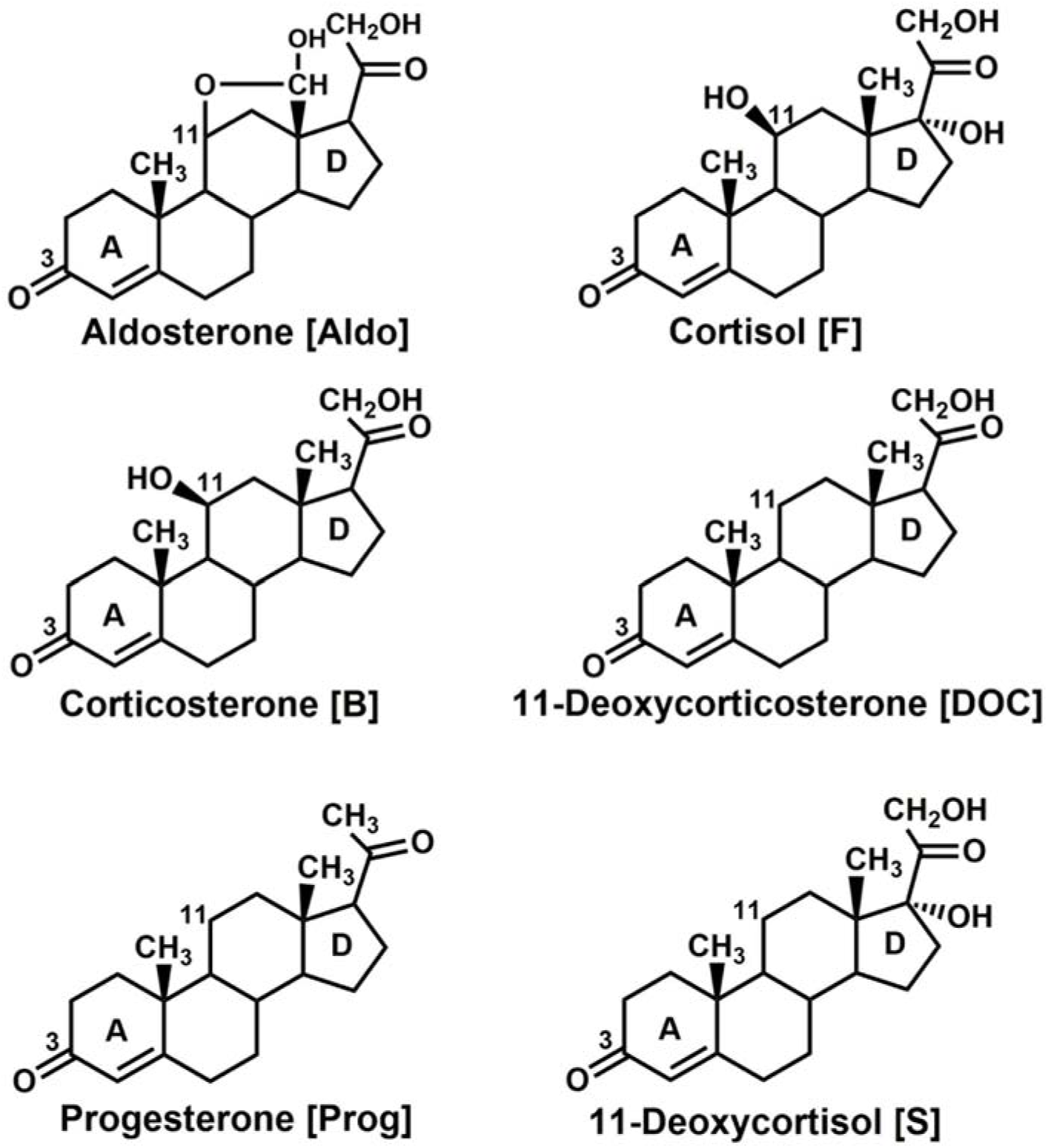
Structures of Mineralocorticoids and Glucocorticoids. Aldosterone [Aldo] and 11-deoxycorticosterone [DOC] are the main physiological mineralocorticoids in vertebrates. Cortisol [F] and corticosterone [B] are the main physiological glucocorticoids in vertebrates. 11-deoxycortisol [S] has activity as mineralocorticoid and glucocorticoid in lamprey CR (Baker 2011). Progesterone [Prog] is an antagonist for human MR (Rupprecht et al. 1993,Geller, 2000 #148)and an agonist for fish MRs (Pippal et al. 2011; Sturm et al. 2005; Sugimoto et al. 2016).

The similar affinity of F, B and Aldo for the MR raised the question of how Aldo could occupy the MR in the presence of F and B, which have from 100 to 1000 fold higher concentrations in human and rodent serum, respectively, than that of Aldo. Human MR should be occupied by F and rodent MR by B. Selective binding of Aldo to the MR in the presence of either F or B arises from a novel enzymatic mechanism involving expression of 11β-hydroxysteroid dehydrogenase-type 2 (11βHSD2) in epithelia containing the MR. This enzyme metabolizes the 11β-OH of F and B to a ketone, yielding cortisone and 11-dehydrocorticosterone, respectively, two inactive steroids (Baker 2010; Chapman, et al. 2013; Draper and Stewart 2005; Edwards, et al. 1988; Funder, et al. 1988; Odermatt and Kratschmar 2012). Aldo is inert to 11βHSD2 and can occupy the kidney and colon MRs in the presence of 11βHSD2. However DOC, S and Prog lack an 11β-OH, which allows these steroids to compete with Aldo for the MR.

Specificity of human MR for Aldo compared to other corticosteroids also comes from Aldo’s strong transcriptional activation of the MR. Indeed, a follow-up paper (Arriza, et al. 1988) showed that the half-maximal response (EC50) of Aldo for human MR was over 100X lower than that of F. Subsequent reports extended the stronger response of the human MR to Aldo compared to other corticosteroids (Hellal-Levy, et al. 1999; Lombes, et al. 1994; Mani, et al. 2016; Rupprecht, et al. 1993; Sugimoto, et al. 2016). Interestingly, Prog is an antagonist for human MR (Geller, et al. 2000; Rupprecht et al. 1993; Sugimoto et al. 2016).

Important in understanding MR evolution is the pathway for the synthesis of corticosteroids that are ligands for the MR (Figure 3). Progressive modification of Prog yields steroids with substituents that are ligands for either the MR and GR or both. The position of each steroid in this pathway appears to coincide with the evolution of steroids in vertebrates as physiological activators of the CR, MR and GR. For example, Aldo, which is at the end of one synthetic pathway, is not present in either lamprey or hagfish serum (Bridgham et al. 2006). Potential activators of the CR are S, DOC and Prog, which have been found in Atlantic sea lamprey serum (Close, et al. 2010; Roberts, et al. 2014; Wang, et al. 2016). These steroids are at the beginning of the pathway. S has mineralocorticoid activity in lamprey (Close et al. 2010); the roles of DOC, which is a mineralocorticoid in mammals (Hawkins, et al. 2012; Lam, et al. 2006) and of Prog are not known.

**Figure 3.**
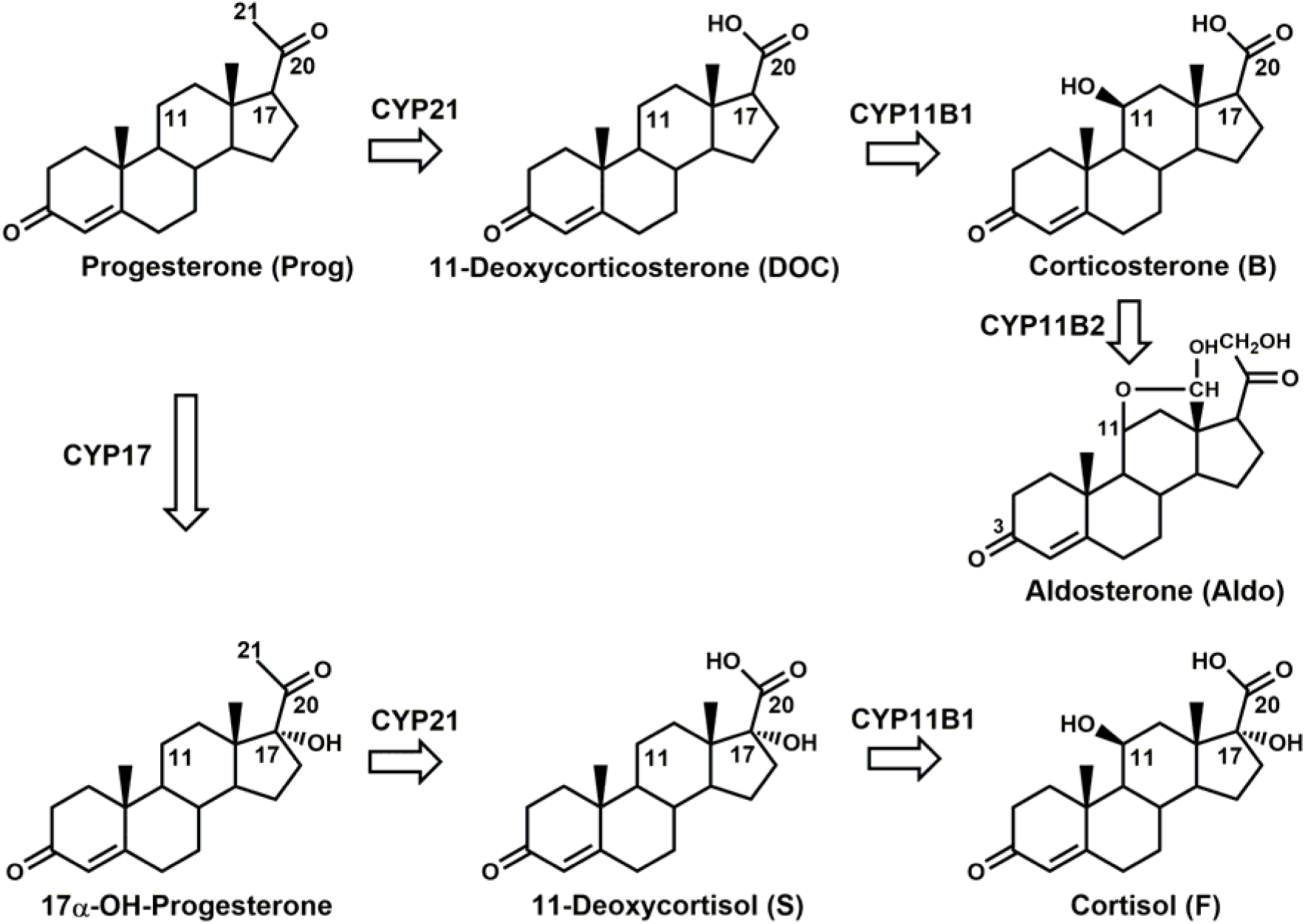
Pathway for synthesis of Aldosterone from Progesterone and Cortisol from 17α-OH-Progesterone. Progesterone is hydroxylated at C21 to form 11-deoxycorticosterone, which is hydroxylated at C11 to form corticosterone. Hydroxylated of corticosterone at C18 followed by oxidation of the C18 hydroxyl forms aldosterone. In a second pathway, progesterone is hydroxylated at C17 and then hydroxylated at C21 to form 11-deoxycortisol, which is hydroxylated at C11 to form cortisol (Baker 2011; Rossier et al. 2015).

Chondrichthyes contain B and a novel derivative 1α-OH-B, which has not been found in other vertebrates. F and B, but not Aldo, are found in ray-finned fish (Jiang, et al. 1998; Sakamoto, et al. 2011). Aldo first appears in tetrapods (Joss, et al. 1994; Rossier, et al. 2015), which also contain F and B.

Interestingly, Aldo has low EC50s for the MR in vertebrates that do not synthesize Aldo. The EC50 of Aldo for hagfish CR is 0.4 nM (Bridgham et al. 2006). The EC50s of Aldo, DOC and B for skate MR are 0.07 nM, 0.03 nM and 0.09 nM, respectively (Carroll et al. 2008). 11βHSD2 first appears in Chondrichthyes (Baker et al. 2015; Rossier et al. 2015), in which 11βHSD2 may regulate access of B to the MR. Although Aldo is not found in ray-finned fish, Aldo is a strong transcriptional activator of fish MR (Greenwood, et al. 2003; Pippal, et al. 2011; Stolte, et al. 2008; Sturm, et al. 2005; Sugimoto et al. 2016). Indeed, the physiological mineralocorticoid in ray-finned fish is not fully understood because several 3-ketosteroids, including F, DOC, B, S and Prog are transcriptional activators of the MR (Greenwood et al. 2003; Pippal et al. 2011; Stolte et al. 2008; Sturm et al. 2005; Sugimoto et al. 2016), and one or more of these steroids could be a physiological mineralocorticoid.

Thus, during the evolution of the MR in cartilaginous fishes, ray-finned fishes and terrestrial vertebrates, there have been changes in MR specificity for corticosteroids, as well as the MR’s physiological function (Hawkins et al. 2012; Jaisser and Farman 2016; Martinerie, et al. 2013; Mifsud and Reul 2016; Rossier et al. 2015; Sakamoto, et al. 2016; Vize and Smith 2004). To gain a deeper understanding of the evolution of the MR, we have taken advantage of the sequencing, in the last five years, of a cornucopia of genomes from vertebrates at key evolutionary transitions, including lamprey, elephant shark and coelacanth, to investigate regions of conservation and divergence among and between MRs and GRs. We use these sequence analyses, the crystal structures human MR (Bledsoe, et al. 2005; Edman, et al. 2015; Fagart, et al. 2005; Li, et al. 2005) and GR (Bledsoe, et al. 2002; He, et al. 2014) and functional studies of human MR mutants (Bledsoe et al. 2005; Fagart, et al. 1998; Geller et al. 2000; Jimenez-Canino, et al. 2016; Li et al. 2005; Shibata, et al. 2013,Mani, 2016 #272) to investigate the evolution of selectivity for Aldo and other corticosteroids by the MR in terrestrial vertebrates (Baker et al. 2013) and ray-finned fish (Arterbery, et al. 2011; Bury and Sturm 2007; Prunet, et al. 2006; Sugimoto et al. 2016), as well as the basis for binding of some glucocorticoids by human MR and other vertebrate MRs (Li et al. 2005; Mani et al. 2016). Together these data provide molecular and structural fingerprints for investigating the evolution of selectivity for 3-ketosteroids by the MR in vertebrates.

### The MR is a multi-domain transcription factor

Like other steroid receptors, the MR is composed of several functional domains (Figure 4). The MR contains an A/B domain at the N-terminus (NTD), a DNA-binding domain (DBD) (C domain) near the center, a short hinge domain (D domain) and a steroid-binding domain (LBD) (E domain) at the C-terminus (Arriza et al. 1988; Huang et al. 2010; Huyet, et al. 2012; Pascual-Le Tallec and Lombes; Yang and Young 2009). The A/B domain contains an activation function 1 [AF1] and the E domain contains an AF2 domain (Faresse 2014; Huyet et al. 2012; Li et al. 2005; Pascual-Le Tallec and Lombes 2005). Each domain in MR is important for transcriptional responses (Faresse 2014; Fuller, et al. 2012).

**Figure 4.**
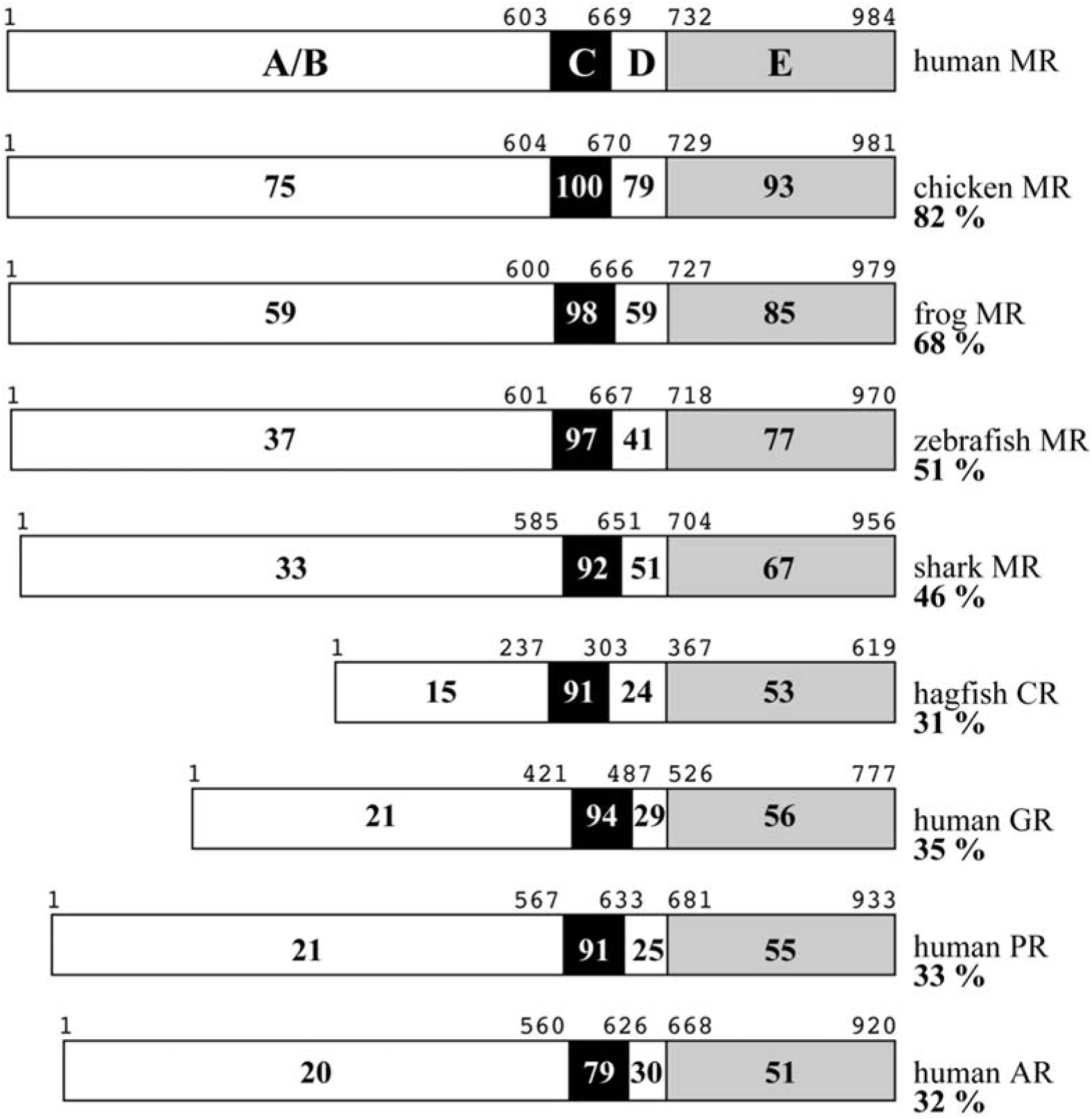
Comparison of Domains on MR, GR, PR, AR and CR. The A/B domain to E domains are schematically represented with the numbers of amino acid residues and the percentage of amino acid identity between the domains compared to human MR. For example, the entire human MR sequence is 82% identical to that of chicken MR, while domain E (LBD) on human MR is 93% identical to that of chicken MR. Accessions are in Supplement Table 1.

As shown in Figure 4, the DBD is highly conserved in vertebrate MRs, while the NTD and hinge domains are poorly conserved. The LBD has intermediate sequence conservation, which makes it useful for phylogenetic analysis of the MR (Baker et al. 2015; Bridgham, et al. 2008; Rossier et al. 2015). In addition, the sequence of the LBD in vertebrate MRs is of interest because mutations in the LBD have been correlated with changes in transcriptional activation by Aldo and other steroids (Bledsoe et al. 2005; Bridgham et al. 2006; Fagart et al. 2005; Fagart et al. 1998; Funder 2013; Geller et al. 2000; Li et al. 2005; Shibata et al. 2013,Mani, 2016 #272). Thus, a multiple alignment of the LBD of key vertebrate MRs and GRs can be used for a phylogenetic analysis of the MR, as well as to identify sites that could be important in functional divergence of vertebrate MRs from each other and from the GR.

### Evolution of vertebrate MR steroid-binding domain: Divergence from its GR paralog

Figure 5 shows a multiple alignment of the steroid-binding domain on various MRs, GRs, CRs, PRs and AR from vertebrates at key evolutionary transitions. Figure 5 also shows the α-helices on the MR as determined by X-ray crystallography (Bledsoe et al. 2005; Edman et al. 2015; Fagart et al. 2005; Li et al. 2005) and amino acids that have been found to be important in either steroid binding or transcriptional activation of the MR or CR (Bledsoe et al. 2005; Bridgham et al. 2006; Edman et al. 2015; Fagart et al. 1998; Geller et al. 2000; Hultman, et al. 2005; Jimenez-Canino et al. 2016; Li et al. 2005; Mani et al. 2016; Shibata et al. 2013) or the GR (Bledsoe et al. 2002; Edman et al. 2015; He et al. 2014; Mani et al. 2016). A striking feature, discussed in more detail below, is the strong conservation of amino acids among the CR, MR, GR, PR and AR, including lamprey PR and CR. Indeed, some amino acids are conserved in vertebrate MRs, GRs and CRs and even in lamprey and hagfish PRs.

**Figure 5.**
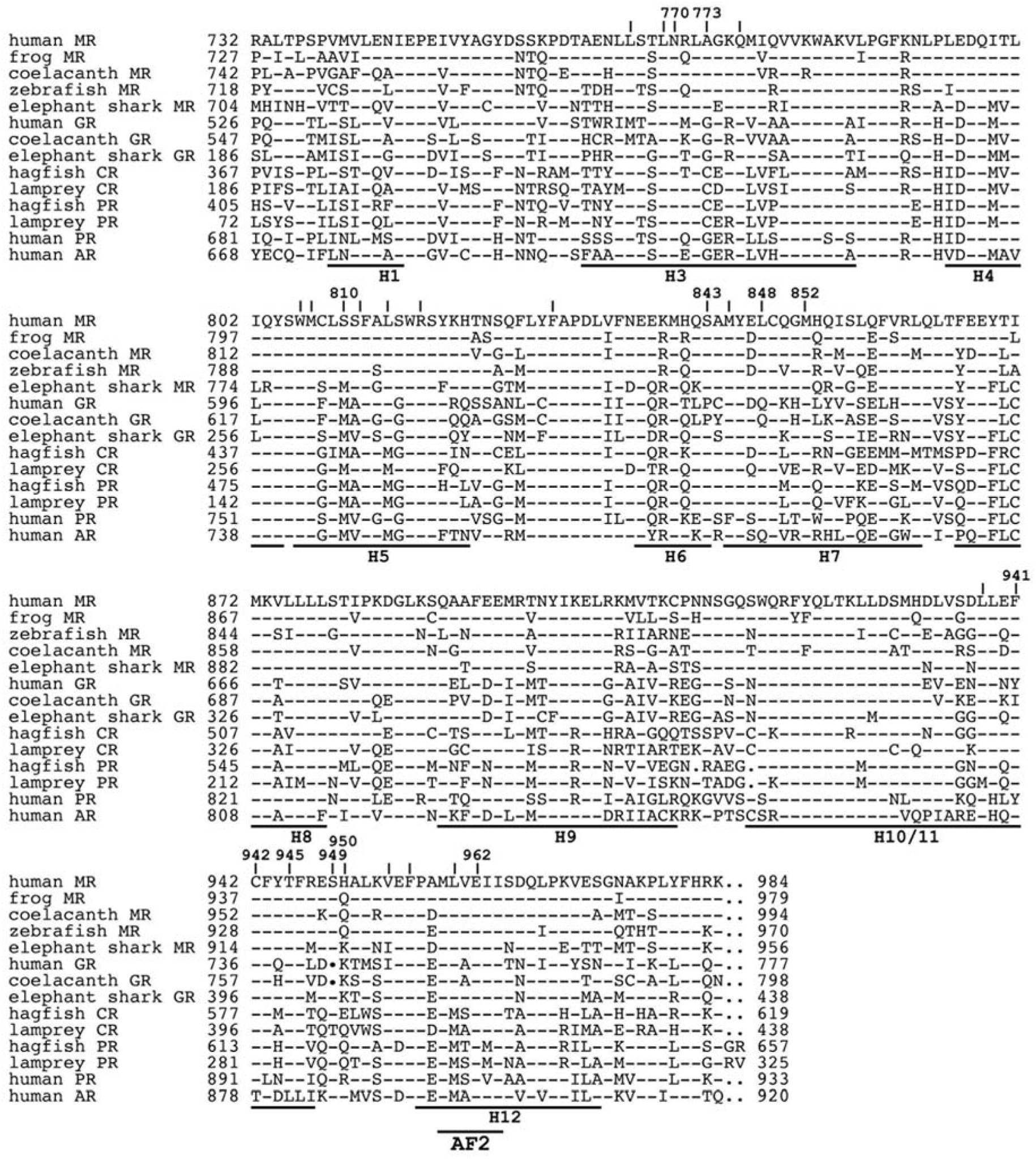
Alignment of the steroid binding domain on vertebrate MRs, CRs, GRs, PRs and AR. The steroid binding domains were collected with BLAST searches of GenBank. Clustal W was used to construct the multiple alignment (Larkin, et al. 2007). The crystal structure of human MR (PDB: 2A3I) {Li, 2005 #10} was use to locate α-helices. Amino acids that contact Aldo are shown above human MR. The highly conserved Glu-962 is part of AF2, which contacts co-activator proteins. The functions of Ser-949 and His-950, remain to be elucidated.

We used this multiple alignment to construct an up-dated phylogeny of the steroid-binding domain on the MR and other 3-ketosteroid receptors (Figure 6). This phylogeny indicates that the CR and PR evolved from an ancestral 3-ketosteroid receptor through gene duplication and divergence (Baker et al. 2015; Rossier et al. 2015; Thornton 2001); that the MR and GR evolved from an ancestral CR; that the MR is closer than the GR is to the CR, and that the CR ancestor of the MR and GR appears to be lost in Pacific and Atlantic lamprey. Also, the presence of the PR and the absence of an AR in cyclostomes indicates that the AR evolved from a duplication of an ancestral PR.

**Figure 6.**
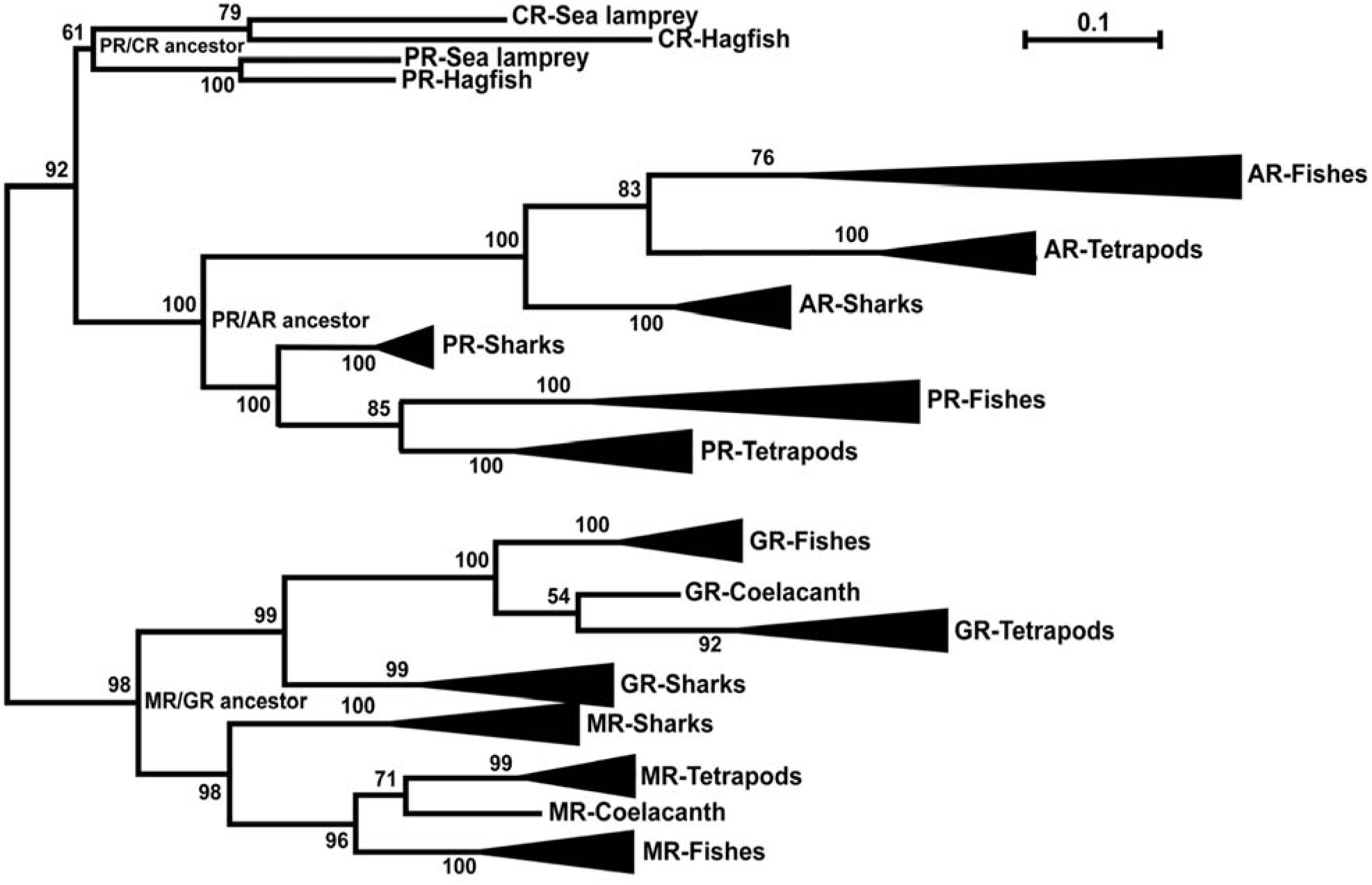
Phylogenetic analysis of vertebrate MRs, GRs, CRs, PRs and ARs. Steroid binding domains were collected with BLAST searches of GenBank. Then Clustal W (Larkin et al. 2007) was used to construct the multiple alignment, and phylogenetic trees were constructed with Maximum likelihood (ML) analysis conducted using JTT+G+I model (Guindon and Gascuel 2003). Statistical confidence for each branch in the tree was evaluated with 1,000 bootstrap runs. MEGA5 program was used for these analysis (Tamura, et al. 2011). A gene duplication of an ancestral 3-ketosteroid receptor in cyclostomes produced the ancestors of the CR and PR in modern lampreys and hagfishes. Lamprey and hagfish CR and lamprey PR cluster in one branch. The MR and GR are in another branch. The CR ancestor of the MR and GR appears to be lost in lamprey and hagfish. In Gnathostomes, a gene duplication produced the AR and PR.

### Evolution of contacts between the MR and A and B rings on Aldo and other 3-ketosteroids

Stabilizing interactions between α-helix 3 and α-helix 5 with each other and with the A and B rings on corticosteroids are important in transcriptional activation of the MR (Baker et al. 2013; Bledsoe et al. 2005; Fuller et al. 2012; Geller et al. 2000; Huyet et al. 2012; Li et al. 2005), as well as other 3-ketosteroid receptors. Consistent with the common ancestry of the MR, GR, PR, AR and structural similarities of the A and B rings in their canonical ligands, some key amino acids in the MR are conserved in the GR, PR, AR and CR (Baker et al. 2013; Li et al. 2005; Mani et al. 2016). However, other amino acids are not conserved, providing specificity for mineralocorticoids (Baker et al. 2007; Baker et al. 2013; Bledsoe et al. 2005; Huyet et al. 2012), glucocorticoids (Bledsoe et al. 2002; He et al. 2014), progestins (Williams and Sigler 1998) and androgens (Sack, et al. 2001) in their cognate receptors.

For example, in human MR, Gln-776 (helix 3) and Arg-817 (helix 5) are conserved in corresponding positions in vertebrate MR, GR, PR, AR and CR (Figure 5, Figure 7). Also conserved in human MR, lamprey CR, as well as the GR, PR and AR are contacts between the side chain on Phe-829 (human MR) with the A ring on corticosteroids and between the backbone oxygen on Phe-829 with Nε and Nη2 on Arg-817 (Figures 5 and 7).

**Figure 7.**
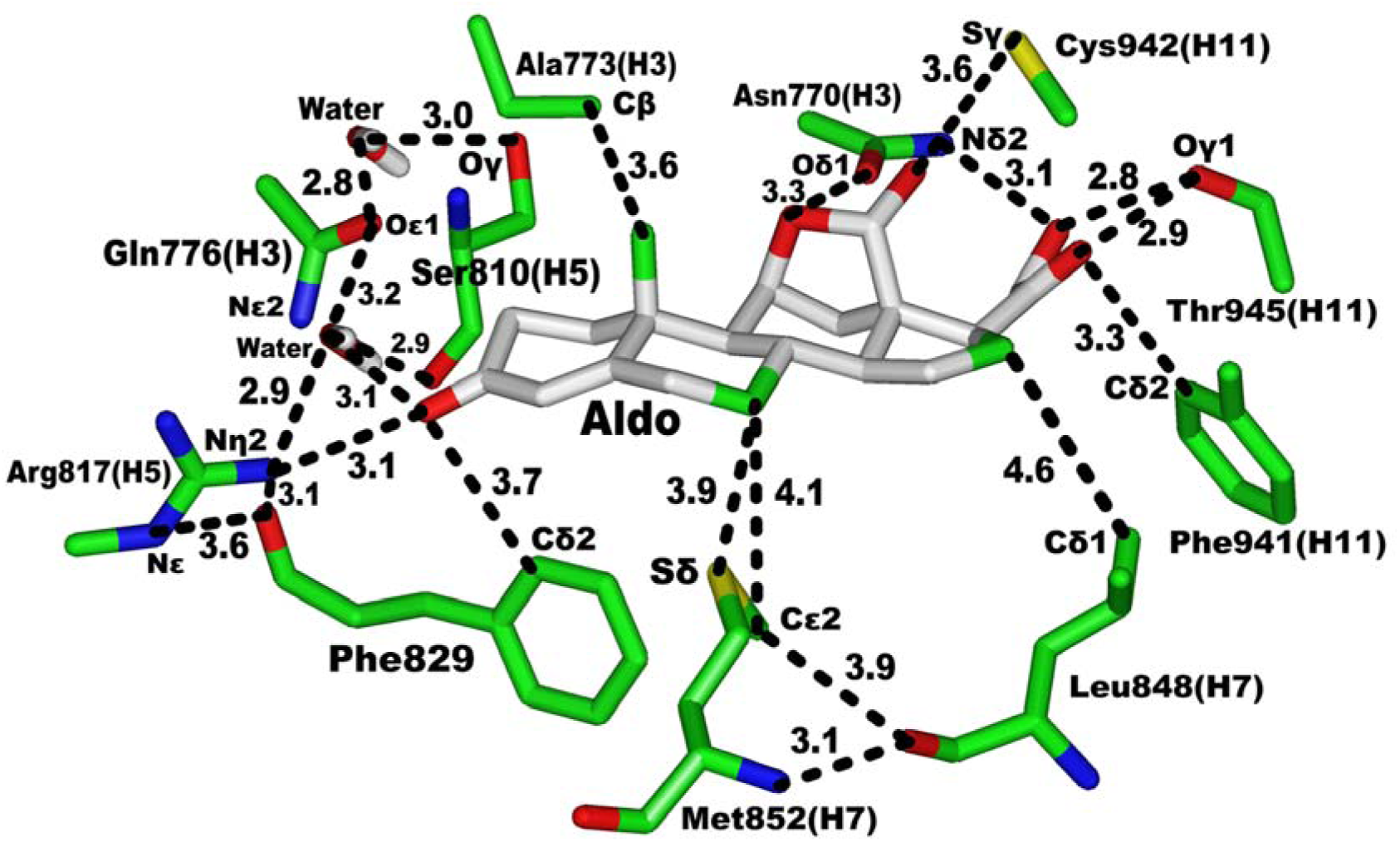
Contacts between the MR and Aldo and two water molecules. Contacts between Aldo with human MR (PDB: 2AA2) (Bledsoe et al. 2005) are shown. Two water molecules mediate contacts between Ser-810 and the A ring of Aldo.

### Ser-810 (helix 5) in human MR: Evolution in the common ancestor of ray-finned and lobe-finned fish

Ser-810 in human MR also is important in binding of the A ring of steroids. The crystal structure of human MR with Aldo reveals that Ser-810 stabilizes the A ring on Aldo through a hydrogen bond network with two water molecules (Bledsoe et al. 2005). In one hydrogen bond network, a water molecule contacts Oγ on Ser-810 and the C3-ketone on Aldo; in another network, a water molecule contacts the backbone oxygen on Ser-810, Nη2 on Arg-817 and the C3-ketone on Aldo (Bledsoe et al. 2005) (Figure 7).

A serine corresponding to Ser-810 in human MR first appears in ray-finned fish and lobe-finned fish (Figure 5) (Baker et al. 2007; Baker et al. 2013; Baker, et al. 2011). In contrast, chondrichthian MRs and cyclostome CRs contain a methionine corresponding to Ser-810. Moreover, the GR, PR, and AR also have a methionine at this position (Figure 5) (Baker et al. 2007; Baker et al. 2013; Baker et al. 2011). Thus, this water-mediated hydrogen bond between Oγ on Ser-810 and C3-ketone on Aldo, which emerged in the MR in common ancestor of ray-finned fish and tetrapods, is unique among 3-ketosteroid receptors. The evolution of this serine in the MR affects binding to 3-ketosteroids because, as noted by Bledsoe et al. (Bledsoe et al. 2005), methionine at this position cannot participate in a water-mediated hydrogen bond with the C3-ketone on corticosteroids, indicating that there was a change in the mechanism for stabilization of the C3-ketone in the MR in ray-finned fish and tetrapods.

Moreover, as discussed below, it appears that replacement of methionine with serine was important the loss transcriptional activation of the MR by Prog (Geller et al. 2000) and cortisone (Rafestin-Oblin, et al. 2003).

### Evolution of the contact between Ser810 (Helix 5)-Ala773 (Helix 3) in human MR: Role in the divergence of the MR and GR

Important evidence for a physiological role of Ser-810 in human MR comes from a report in 2000 by Geller et al. (Geller et al. 2000), who identified a Ser810Leu mutation in the MR, which was activated by Prog (EC50 of ~ 1nM). Prog activation of the MR is unexpected because Prog is an antagonist for wild-type human MR (Geller et al. 2000; Rafestin-Oblin et al. 2003; Rupprecht et al. 1993; Sugimoto et al. 2016). The mineralocorticoid activity of Prog for Leu-810 MR explained high blood pressure in pregnant woman with this mutant MR. In addition, cortisone, which binds poorly to human MR, is an agonist for the Leu-810 MR (Rafestin-Oblin et al. 2003) and could cause hypertension in people with this mutant MR. Moreover, spironolactone, an MR antagonist, activated the Ser810Leu MR in COS-7 cells. Thus, the evolution of an ancestral Ser-810 in the MR in ray-finned fish and tetrapods has an important physiological consequence in preventing activation of the MR by Prog and cortisone.

A 3D model of Leu810-MR found a van der Waals contact between Leu-810 and Ala-773 in the mutant MR, which stabilized the contact between helix 3 and helix 5 (Geller et al. 2000). Transcriptional analyses of MRs with mutations at 810 and 773 supported stabilization of the helix 3-helix 5 contact in the agonist activity of Prog. Subsequent crystal structures of Leu810MR found a stabilizing interaction between helix 3 and helix 5 (Bledsoe et al. 2005; Fagart et al. 2005). This contact between Ala-773 and Ser-810 is not found in crystal structure of wild-type human MR (Bledsoe et al. 2005; Li et al. 2005).

As mentioned previously, Ser-810 evolved in the MR in ray-finned fish and tetrapods. Lamprey and hagfish CR have cysteine (Cys-227) and methionine (Met-264) corresponding to Ala-773 and Ser-810, respectively. A 3D model of lamprey CR found a van der Waals contact between Cys-227 and Met-264. In skate and elephant shark MR, this cysteine is replaced with the alanine that is conserved MR descendants. Based on mutagenesis studies of Geller et al. (Geller et al. 2000), we predict that Ala-191 and Met-238 in skate MR and Ala-745 and Met-782 in elephant shark MR (corresponding to human MR Ser-810), will have van der Waals contacts and, thus, Prog will be an agonist for skate and elephant shark MRs.

The evolution of this helix 3-helix 5 contact in the GR affects its response to 3-ketosteroids and the divergence of the GR and MR. Gly-106 in skate GR and Gly-227 in elephant shark GR correspond to Ala-191 in skate MR and Ala-745 in elephant shark MR. The GR in tetrapods and ray-finned fish conserves a corresponding glycine (helix 3) and methionine (helix 5). The human GR crystal structure (Bledsoe et al. 2002; Zhang, et al. 2005) reveals, as expected, that Gly-567 (helix 3), which lacks a side chain, does not contact Met-604 (helix 5). Interestingly, replacement of Gly-567 with Ala-567 decreases the response to F, B and DEX by at least 10-fold (Zhang et al. 2005). In skate and elephant shark MR, the corresponding site contains an alanine suggesting that the emergence in cartilaginous fish GRs of a glycine corresponding to Gly-567 in human GR was important in evolution of specificity for glucocorticoids.

### Evolution of contacts between the MR and C and D rings on 3-ketosteroids

Crystal structures of the MR reveal that differences in contacts between the MR and hydroxyl groups on the C and D rings of 3-ketosteroids (Figure 2) influence their transcriptional activity for the various MRs, as well as for the GR and other steroid receptors (Bledsoe et al. 2005; Bledsoe et al. 2002; Huang et al. 2010; Huyet et al. 2012).

Vertebrate MRs, CRs and chondrichthian GR and PR conserve many amino acids in human MR (Figure 5) that contact the C and D rings on Aldo (Figure 7), DOC and B. These include Asn-770 (helix 3), Met-852 (helix 7), Phe-941 (helix 11), Cys-942 (helix 11) and Thr-945 (helix 11) (Figure 5). Tetrapod and ray-finned fish GRs also conserve amino acids corresponding to Asn-770, Met-852, Cys-942 and Thr-945, but not Phe-941 in human MR. Interestingly lamprey PR conserves amino acids corresponding to Asn-770, Met-852, Phe-941, Cys-942 and Thr-945 in human MR.

### Ser-843 (helix 6) in human MR: Role in divergence from the GR

Analysis of the crystal structure of the human MR with B (Li et al. 2005) and human GR with DEX (Bledsoe et al. 2002) identified a pocket containing helices 6 and 7 that was present in the GR and not in the MR. This pocket on the GR could accommodate a 17α-hydroxyl group on F and DEX and glucocorticoids with other 17α substituents. Two amino acid differences between human MR and GR (Ser-843 and Leu-848 in human MR, Pro-637 and Gln-642 in human GR) (Figure 5) were identified as important in this conformational change. Indeed, when human GR and MR are superimposed, Ser-843 in the MR is displaced by over 5 Å from Pro-637 in the GR and Leu-848 is 4.5 Å from C16 on B (Baker et al. 2013; Rossier et al. 2015) (Figure 8), which could be important in different responses between MR and GR to F and DEX (Li et al. 2005). In human GR, Gln-642 has a hydrogen bond with the 17α-hydroxyl on DEX (Figure 8) (Bledsoe et al. 2002; He et al. 2014). In the MR, the hydrophobic side chain on Leu-848 was proposed to clash with the 17α-hydroxyl on F and DEX. In contrast, B, Aldo and DOC, which lack a 17α-hydroxyl, would not clash with Leu-848, explaining the stronger response of the MR to these steroids. However, a crystal structure of the MR with DEX (Edman et al. 2015) did not find a steric clash between Leu-848 on the MR with the 17α-hydroxyl on Dex suggesting that other sites on the MR and GR are important in their transcriptional response to F and other corticosteroids with a 17α-hydroxyl group.

**Figure 8.**
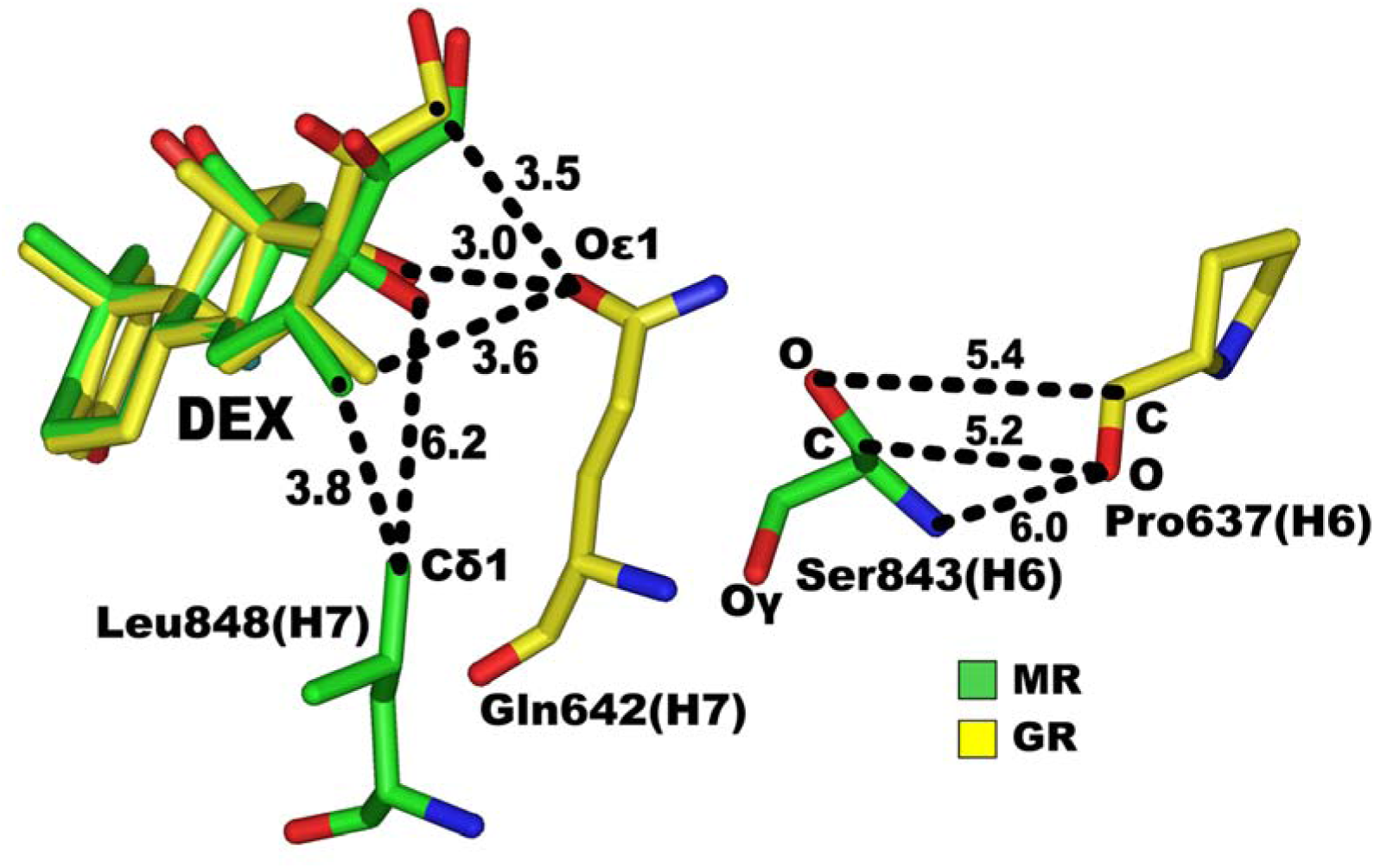
Comparison of Ser-843/Leu-848 on human MR with Pro-637/Gln-642 on human GR. Human GR complexed with dexamethasone (PDB: 1M2Z) (Bledsoe et al. 2002) and human MR complexed with dexamethasone (PDB: 4UDA) (Edman et al. 2015) were superimposed. In human MR, Cδ1 on Leu-848 is 6.2 Å and 3.8 Å, respectively, from 17α-OH and 16α-CH3 on DEX. In human GR, Oε1 is 3.0 Å and 3.6 Å, respectively, from 17α-OH and 16α-CH3 on DEX. Ser-843 and Pro-637 are displaced by over 5 Å.

Supporting this hypothesis are the low EC50s of F for fish MR, which conserve a serine and leucine corresponding to Ser-843 and Leu-848 in human MR. The EC50 of F is 0.02 nM for cichlid (Greenwood et al. 2003), 1.1 nM for trout (Sturm et al. 2005), 2.4 nM for carp (Stolte et al. 2008) and 0.22 nM for zebrafish (Pippal et al. 2011).

Nevertheless, the Ser-Pro mutation in helix 6 in the MR likely has some biochemical effect that is important in divergence of the GRs in ray-finned fish and terrestrial vertebrates from the GR and MR in cartilaginous fish. Indeed, mutagenesis of amino acids in an ancestral CR (AncCR) corresponding to Ser-843 and Leu-848 was incorporated into a novel model to investigate the evolution specificity for steroids with a 17α-hydroxyls such as F for the GR (Bridgham et al. 2006). First, AncCR was transfected into cells and exposed to Aldo or F. The AncCR had a strong response to Aldo and a weak response to F. Then Ser-106 and Leu-111 on AncCR, corresponding to Ser-843 and Leu-848, were mutated to Pro and Gln, as found in ray-finned fish and terrestrial vertebrate GRs. The AncCR-Gln111 mutant had low activity for Aldo, F and DOC, while AncCR-Pro106 was activated by Aldo, DOC and F. The subsequent double AncCR-Pro106/Gln111 mutant had an increased response to F and low response to Aldo, indicating that the GR evolved from AncCR through a step wise mutation of Ser-106 to Pro followed by Leu-111 to Gln. However, studies with human MR (Li et al. 2005; Mani et al. 2016) find that Leu843Gln human MR mutant has a favorable response to F, unlike that of the AncCR, leaving unresolved the pathway for the formation of Pro and Gln in the GR. Future studies with mutations at the corresponding serine and leucine residues in GRs and MRs in cartilaginous fish should provide more direct data on the pathway for the evolution of specificity for corticosteroids in the GR and MR.

### Phosphorylation of Ser-843 inactivates human MR

An important discovery of another physiological role of Ser-843, also relevant for the Ser to Pro mutation in the GR ancestor of lobe-finned and ray-finned fish, comes from a report by Shibata et al. (Shibata et al. 2013) showing that under normal conditions Ser-843 in human MR is phosphorylated in intercalated cells in the kidney distal tubule and this phosphorylated MR is inactive. De-phosphorylation of Ser-843 by a phosphatase induced by angiotensin II activates the MR, such that binding of Aldo leads to sodium chloride absorption and potassium secretion. Interestingly, high potassium levels increase phosphorylation of Ser-843 (Funder 2013; Jimenez-Canino et al. 2016; Shibata et al. 2013). Phosphorylated Ser-843 human MR has only been found in intercalated cells in the kidney distal tubule; other cells do not contain phosphorylated MR.

A serine corresponding to Ser-843 is found in lamprey and hagfish CR, cartilaginous fish MRs and GRs and lamprey PR. This serine also is conserved in descendent MRs, PRs and ARs. If skate MR Ser-261 and GR Ser-176 and elephant shark MR Ser-815 and GR Ser-297, are phosphorylated *in vivo*, then the evolution of a corresponding proline in the GR lobe-finned and ray-finned fishes would provide a mechanism for specificity for regulation of transcriptional activation of the MR through a kinase/phosphatase that would not affect the GR.

At this time, it is not known if this serine is phosphorylated in lamprey PR or other PRs and ARs, or if phosphorylation alters the response to steroids.

### Unanswered questions

Dobzhansky’s aphorism “Nothing in Biology Makes Sense Except in the Light of Evolution” (Dobzhansky.T 1973) is our lodestar for investigating the evolution of the MR as well as other steroid receptors and steroidogenic enzymes. In this spirit we discuss other properties of the MR that merit further investigation to shed light on the evolution of the MR.

### Transcriptional activation of fish MR by Prog, a possible mineralocorticoid

The absence of Aldo in fish has led to speculation that F and DOC may be a physiological mineralocorticoid for fish (Arterbery et al. 2011; Baker 2003; Baker et al. 2007; Bury and Sturm 2007; McCormick, et al. 2008; Prunet et al. 2006; Sakamoto et al. 2011; Sakamoto et al. 2016; Sturm et al. 2005; Takahashi and Sakamoto 2013). Interestingly Prog is a transcriptional activator of ray-finned fish, which also respond to 19-norProg and spironolactone (Pippal et al. 2011; Sturm et al. 2005; Sugimoto et al. 2016). This response is unexpected because Ser-810 in human MR is crucial for the absence of transcriptional activation by Prog, spironolactone and 19-norProg (Baker et al. 2013; Bledsoe et al. 2005; Fagart et al. 2005; Geller et al. 2000), and fish MR contain a serine corresponding to Ser-810 in human MR. The basis for this novel response to Prog, spironolactone and 19-norProg is not known. Nevertheless, Prog may be a physiological mineralocorticoid in fish. Like DOC, Prog lacks an 11β-hydroxyl and thus is inert to 11β-HSD2 (Chapman et al. 2013; Odermatt and Kratschmar 2012).

### Function of Ser-849 in human MR and its deletion in tetrapod and ray-finned fish GRs

Human MR contains Ser-949 in the loop connecting helix 11 and helix 12. A corresponding serine is found in other MRs, shark GR, the CR, lamprey and human PR and human AR, but not in the GR in tetrapods and ray-finned fish [Figure 5] (Baker et al. 2007; Baker et al. 2013). The physiological consequences of this serine in human MR and its deletion in the GR are not known. This difference between the GR and MR appears to alter the conformation of helix 12, which contains AF2, in human GR and MR [Figure 9]. Differences in the conformation of AF2 may be important in selective binding of co-activators to MR and GR (Fuller et al. 2012; Hu and Funder 2006; Hultman et al. 2005; Li et al. 2005; Yang and Young 2009).

**Figure 9.**
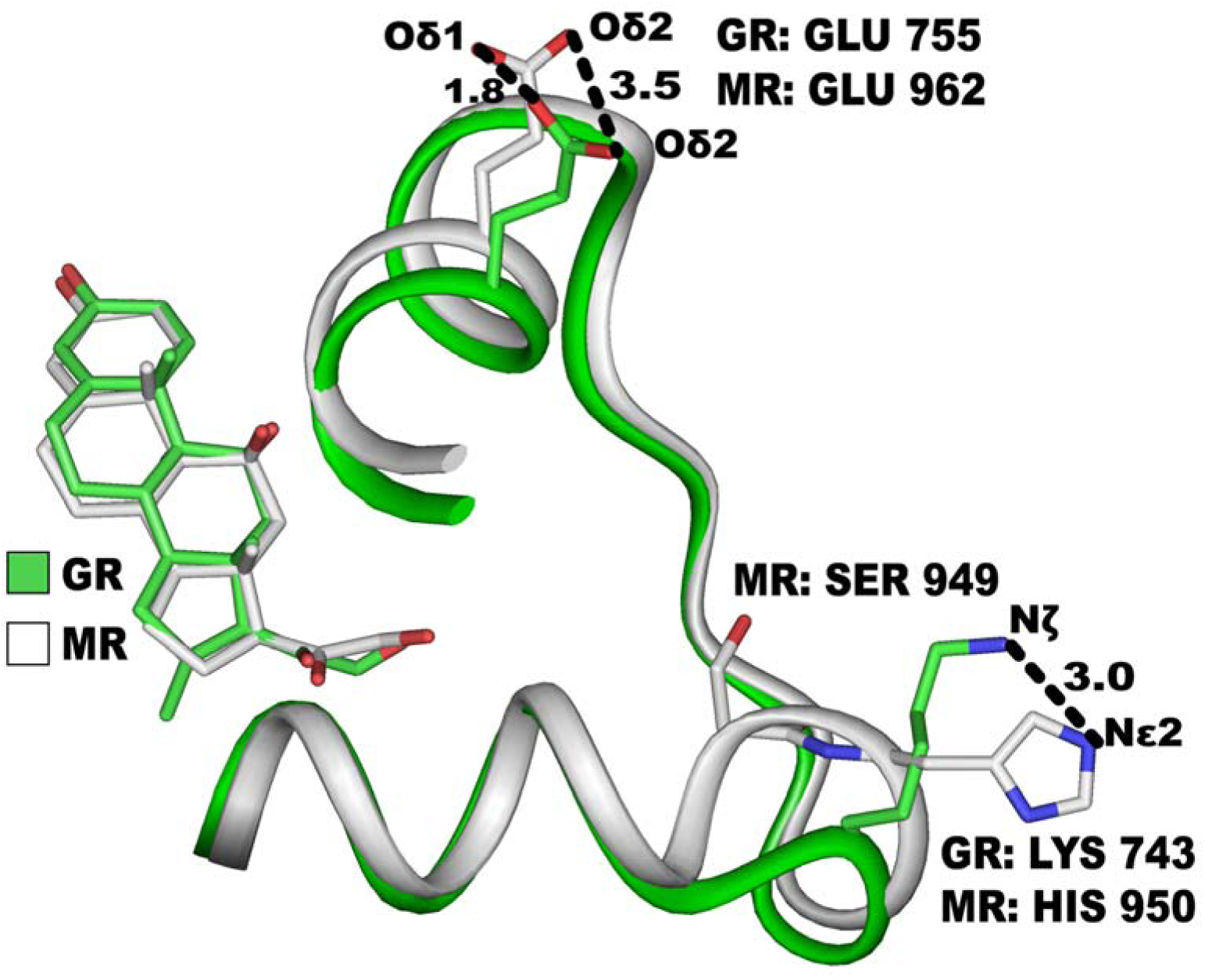
Superposition of helix 12 and the preceding loop in human MR and human GR. Human GR has a deletion corresponding to Ser-949 in human MR, which is in the loop connecting helix 11 and helix 12 in human MR. In human GR, this deletion displaces Oδ2 on Glu-755 in helix 12 on human GR from Oδ2 on Glu-962 in human MR by 3.5 Å. In the loop preceding helix 12, Nζ on Lys-743 on human GR is 3 Å from Nε2 on His-950 on human MR. Glu-755 and Glu-962 in the AF2 domain are highly conserved and are part of charge clamp 1 between co-activators and the GR and MR (Kattoula and Baker 2014; Li et al. 2005).

### His-950 in human MR: role in the evolution in old world monkeys

A histidine corresponding to His-950 evolved in the MR in old world monkeys (Baker et al. 2007), which separated from new world monkeys about 40 million years ago. The MR in new world monkeys, other primates, birds, amphibians, coelacanths and ray-finned fish contains glutamine at this position [Figure 5] (Baker et al. 2013). The functional basis for mutation of a highly conserved glutamine to a histidine, amino acids with different structures, is not known. Nevertheless, the differences between glutamine and histidine suggest that this is not a neutral mutation.

### Transcriptional activation by MR-GR heterodimers

Mammalian MR and GR regulate gene transcription as homodimers (Liu, et al. 1995; Mifsud and Reul 2016). However, reflecting the kinship of the MR and GR, there is evidence that they form functional heterodimers with different properties than their homodimers (Bradbury, et al. 1994; Liu et al. 1995; Ou, et al. 2001; Trapp, et al. 1994). In human hippocampus, stress increases cortisol to levels that occupy the MR and GR. Cortisol activated MR-GR heterodimers bind to glucocorticoid response elements, regulating glucocorticoid target genes (Mifsud and Reul 2016). Recently, heterodimers between trout MR and GR were studied in detail, and the MR in the presence of either cortisol or DOC was found to be a dominant negative repressor of trout GR (Kiilerich, et al. 2015). Thus, the actions of the MR and GR in cells that co-express both receptors is complex and can be influenced by steroids that bind both receptors or are selective for each receptor.

Conservation of functional MR-GR heterodimers over 400 million years suggests that MR-GR heterodimers confer some selective advantage(s) in vertebrates. One possible activity of MR-GR heterodimers in fish comes from cortisol activation of the GR in fish, regulating electrolyte balance (Cruz, et al. 2013; Kumai, et al. 2012). Mineralocorticoid activity of the GR is surprising and raises the question: what role, if any, does the MR have in osmoregulation in fish? One possibility is that fish MR influences osmoregulation through formation of MR-GR heterodimers. In any event, conservation of heterodimer formation between the MR and its GR kin during the evolution of tetrapods and ray-finned fish and suggests new avenues of research to elucidate physiological responses to corticosteroids by the MR and GR.

## Author contributions

M.E.B and J.K. conceived of and wrote the paper.

## References

Arriza JL, Simerly RB, Swanson LW & Evans RM 1988 The neuronal mineralocorticoid receptor as a mediator of glucocorticoid response. Neuron 1 887–900.

Arriza JL, Weinberger C, Cerelli G, Glaser TM, Handelin BL, Housman DE & Evans RM 1987 Cloning of human mineralocorticoid receptor complementary DNA: structural and functional kinship with the glucocorticoid receptor. Science 237 268–275.

Arterbery AS, Fergus DJ, Fogarty EA, Mayberry J, Deitcher DL, Lee Kraus W & Bass AH 2011 Evolution of ligand specificity in vertebrate corticosteroid receptors. BMC Evol Biol 11 14.

Baker ME 2003 Evolution of glucocorticoid and mineralocorticoid responses: go fish. Endocrinology 144 4223–4225.

Baker ME 2010 11Beta-hydroxysteroid dehydrogenase-type 2 evolved from an ancestral 17beta-hydroxysteroid dehydrogenase-type 2. Biochem Biophys Res Commun 399 215–220.

Baker ME 2011 Origin and diversification of steroids: co-evolution of enzymes and nuclear receptors. Mol Cell Endocrinol 334 14–20.

Baker ME, Chandsawangbhuwana C & Ollikainen N 2007 Structural analysis of the evolution of steroid specificity in the mineralocorticoid and glucocorticoid receptors. BMC Evol Biol 7 24.

Baker ME, Funder JW & Kattoula SR 2013 Evolution of hormone selectivity in glucocorticoid and mineralocorticoid receptors. J Steroid Biochem Mol Biol 137 57–70.

Baker ME, Nelson DR & Studer RA 2015 Origin of the response to adrenal and sex steroids: Roles of promiscuity and co-evolution of enzymes and steroid receptors. J Steroid Biochem Mol Biol 151 12–24.

Baker ME, Uh KY & Asnaashari P 2011 3D models of lamprey corticoid receptor complexed with 11-deoxycortisol and deoxycorticosterone. Steroids 76 1451–1457.

Bledsoe RK, Madauss KP, Holt JA, Apolito CJ, Lambert MH, Pearce KH, Stanley TB, Stewart EL, Trump RP, Willson TM, et al. 2005 A ligand-mediated hydrogen bond network required for the activation of the mineralocorticoid receptor. J Biol Chem 280 31283–31293.

Bledsoe RK, Montana VG, Stanley TB, Delves CJ, Apolito CJ, McKee DD, Consler TG, Parks DJ, Stewart EL, Willson TM, et al. 2002 Crystal structure of the glucocorticoid receptor ligand binding domain reveals a novel mode of receptor dimerization and coactivator recognition. Cell 110 93–105.

Bradbury MJ, Akana SF & Dallman MF 1994 Roles of type I and II corticosteroid receptors in regulation of basal activity in the hypothalamo-pituitary-adrenal axis during the diurnal trough and the peak: evidence for a nonadditive effect of combined receptor occupation. Endocrinology 134 1286–1296.

Bridgham JT, Brown JE, Rodriguez-Mari A, Catchen JM & Thornton JW 2008 Evolution of a new function by degenerative mutation in cephalochordate steroid receptors. PLoS Genet 4 e1000191.

Bridgham JT, Carroll SM & Thornton JW 2006 Evolution of hormone-receptor complexity by molecular exploitation. Science 312 97–101.

Bridgham JT, Eick GN, Larroux C, Deshpande K, Harms MJ, Gauthier ME, Ortlund EA, Degnan BM & Thornton JW 2010 Protein evolution by molecular tinkering: diversification of the nuclear receptor superfamily from a ligand-dependent ancestor. PLoS Biol 8.

Bury NR & Sturm A 2007 Evolution of the corticosteroid receptor signalling pathway in fish. Gen Comp Endocrinol 153 47–56.

Carroll SM, Bridgham JT & Thornton JW 2008 Evolution of hormone signaling in elasmobranchs by exploitation of promiscuous receptors. Mol Biol Evol 25 2643–2652.

Chapman K, Holmes M & Seckl J 2013 11beta-hydroxysteroid dehydrogenases: intracellular gate-keepers of tissue glucocorticoid action. Physiol Rev 93 1139–1206.

Close DA, Yun SS, McCormick SD, Wildbill AJ & Li W 2010 11-deoxycortisol is a corticosteroid hormone in the lamprey. Proc Natl Acad Sci U S A 107 13942–13947.

Cruz SA, Lin CH, Chao PL & Hwang PP 2013 Glucocorticoid receptor, but not mineralocorticoid receptor, mediates cortisol regulation of epidermal ionocyte development and ion transport in zebrafish (danio rerio). PLoS One 8 e77997.

Dobzhansky. T 1973 Nothing in Biology Makes Sense except in the Light of Evolution. American Biology Teacher 35 125–129.

Draper N & Stewart PM 2005 11beta-hydroxysteroid dehydrogenase and the pre-receptor regulation of corticosteroid hormone action. The Journal of endocrinology 186 251–271.

Edman K, Hosseini A, Bjursell MK, Aagaard A, Wissler L, Gunnarsson A, Kaminski T, Kohler C, Backstrom S, Jensen TJ, et al. 2015 Ligand Binding Mechanism in Steroid Receptors: From Conserved Plasticity to Differential Evolutionary Constraints. Structure 23 2280–2290.

Edwards CR, Stewart PM, Burt D, Brett L, McIntyre MA, Sutanto WS, de Kloet ER & Monder C 1988 Localisation of 11 beta-hydroxysteroid dehydrogenase--tissue specific protector of the mineralocorticoid receptor. Lancet 2 986–989.

Evans RM 1988 The steroid and thyroid hormone receptor superfamily. Science 240 889–895.

Fagart J, Huyet J, Pinon GM, Rochel M, Mayer C & Rafestin-Oblin ME 2005 Crystal structure of a mutant mineralocorticoid receptor responsible for hypertension. Nat Struct Mol Biol 12 554–555.

Fagart J, Wurtz JM, Souque A, Hellal-Levy C, Moras D & Rafestin-Oblin ME 1998 Antagonism in the human mineralocorticoid receptor. EMBO J 17 3317–3325.

Faresse N 2014 Post-translational modifications of the mineralocorticoid receptor: How to dress the receptor according to the circumstances? J Steroid Biochem Mol Biol 143 334–342.

Fuller PJ, Yao Y, Yang J & Young MJ 2012 Mechanisms of ligand specificity of the mineralocorticoid receptor. J Endocrinol 213 15–24.

Funder JW 2013 Angiotensin retains sodium by dephosphorylating mineralocorticoid receptors in renal intercalated cells. Cell Metab 18 609–610.

Funder JW, Pearce PT, Smith R & Smith AI 1988 Mineralocorticoid action: target tissue specificity is enzyme, not receptor, mediated. Science 242 583–585.

Geller DS, Farhi A, Pinkerton N, Fradley M, Moritz M, Spitzer A, Meinke G, Tsai FT, Sigler PB & Lifton RP 2000 Activating mineralocorticoid receptor mutation in hypertension exacerbated by pregnancy. Science 289 119–123.

Greenwood AK, Butler PC, White RB, DeMarco U, Pearce D & Fernald RD 2003 Multiple corticosteroid receptors in a teleost fish: distinct sequences, expression patterns, and transcriptional activities. Endocrinology 144 4226–4236.

Guindon S & Gascuel O 2003 A simple, fast, and accurate algorithm to estimate large phylogenies by maximum likelihood. Syst Biol 52 696–704.

Hawkins UA, Gomez-Sanchez EP, Gomez-Sanchez CM & Gomez-Sanchez CE 2012 The ubiquitous mineralocorticoid receptor: clinical implications. Curr Hypertens Rep 14 573–580.

He Y, Yi W, Suino-Powell K, Zhou XE, Tolbert WD, Tang X, Yang J, Yang H, Shi J, Hou L, et al. 2014 Structures and mechanism for the design of highly potent glucocorticoids. Cell Res 24 713–726.

Hellal-Levy C, Couette B, Fagart J, Souque A, Gomez-Sanchez C & Rafestin-Oblin M 1999 Specific hydroxylations determine selective corticosteroid recognition by human glucocorticoid and mineralocorticoid receptors. FEBS Lett 464 9–13.

Hu X & Funder JW 2006 The evolution of mineralocorticoid receptors. Mol Endocrinol 20 1471–1478.

Huang P, Chandra V & Rastinejad F 2010 Structural overview of the nuclear receptor superfamily: insights into physiology and therapeutics. Annual review of physiology 72 247–272.

Hultman ML, Krasnoperova NV, Li S, Du S, Xia C, Dietz JD, Lala DS, Welsch DJ & Hu X 2005 The ligand-dependent interaction of mineralocorticoid receptor with coactivator and corepressor peptides suggests multiple activation mechanisms. Mol Endocrinol 19 1460–1473.

Huyet J, Pinon GM, Fay MR, Rafestin-Oblin ME & Fagart J 2012 Structural determinants of ligand binding to the mineralocorticoid receptor. Mol Cell Endocrinol 350 187–195.

Jaisser F & Farman N 2016 Emerging Roles of the Mineralocorticoid Receptor in Pathology: Toward New Paradigms in Clinical Pharmacology. Pharmacol Rev 68 49–75.

Jiang JQ, Young G, Kobayashi T & Nagahama Y 1998 Eel (Anguilla japonica) testis 11beta-hydroxylase gene is expressed in interrenal tissue and its product lacks aldosterone synthesizing activity. Mol Cell Endocrinol 146 207–211.

Jimenez-Canino R, Fernandes MX & Alvarez de la Rosa D 2016 Phosphorylation of Mineralocorticoid Receptor Ligand Binding Domain Impairs Receptor Activation and Has a Dominant Negative Effect over Non-phosphorylated Receptors. J Biol Chem 291 19068–19078.

Joss JMP, Arnoldreed DE & Balment RJ 1994 The Steroidogenic Response to Angiotensin-Ii in the Australian Lungfish, Neoceratodus-Forsteri. Journal of Comparative Physiology B-Biochemical Systemic and Environmental Physiology 164 378–382.

Kattoula SR & Baker ME 2014 Structural and evolutionary analysis of the co-activator binding domain in vertebrate progesterone receptors. J Steroid Biochem Mol Biol 141 7–15.

Kiilerich P, Triqueneaux G, Christensen NM, Trayer V, Terrien X, Lombes M & Prunet P 2015 Interaction between the trout mineralocorticoid and glucocorticoid receptors in vitro. J Mol Endocrinol 55 55–68.

Kumai Y, Nesan D, Vijayan MM & Perry SF 2012 Cortisol regulates Na+ uptake in zebrafish, Danio rerio, larvae via the glucocorticoid receptor. Mol Cell Endocrinol 364 113–125.

Lam EY, Funder JW, Nikolic-Paterson DJ, Fuller PJ & Young MJ 2006 Mineralocorticoid receptor blockade but not steroid withdrawal reverses renal fibrosis in deoxycorticosterone/salt rats. Endocrinology 147 3623–3629.

Larkin MA, Blackshields G, Brown NP, Chenna R, McGettigan PA, McWilliam H, Valentin F, Wallace IM, Wilm A, Lopez R, et al. 2007 Clustal W and Clustal X version 2.0. Bioinformatics 23 2947–2948.

Li Y, Suino K, Daugherty J & Xu HE 2005 Structural and biochemical mechanisms for the specificity of hormone binding and coactivator assembly by mineralocorticoid receptor. Mol Cell 19 367–380.

Liu W, Wang J, Sauter NK & Pearce D 1995 Steroid receptor heterodimerization demonstrated in vitro and in vivo. Proc Natl Acad Sci U S A 92 12480–12484.

Lombes M, Kenouch S, Souque A, Farman N & Rafestin-Oblin ME 1994 The mineralocorticoid receptor discriminates aldosterone from glucocorticoids independently of the 11 beta-hydroxysteroid dehydrogenase. Endocrinology 135 834–840.

Mani O, Nashev LG, Livelo C, Baker ME & Odermatt A 2016 Role of Pro-637 and Gln-642 in human glucocorticoid receptors and Ser-843 and Leu-848 in mineralocorticoid receptors in their differential responses to cortisol and aldosterone. J Steroid Biochem Mol Biol 159 31–40.

Markov GV, Tavares R, Dauphin-Villemant C, Demeneix BA, Baker ME & Laudet V 2009 Independent elaboration of steroid hormone signaling pathways in metazoans. Proc Natl Acad Sci U S A 106 11913–11918.

Martinerie L, Munier M, Le Menuet D, Meduri G, Viengchareun S & Lombes M 2013 The mineralocorticoid signaling pathway throughout development: expression, regulation and pathophysiological implications. Biochimie 95 148–157.

McCormick SD, Regish A, O’Dea MF & Shrimpton JM 2008 Are we missing a mineralocorticoid in teleost fish? Effects of cortisol, deoxycorticosterone and aldosterone on osmoregulation, gill Na+,K+ -ATPase activity and isoform mRNA levels in Atlantic salmon. General and comparative endocrinology 157 35–40.

Mifsud KR & Reul JM 2016 Acute stress enhances heterodimerization and binding of corticosteroid receptors at glucocorticoid target genes in the hippocampus. Proc Natl Acad Sci U S A 113 11336–11341.

Odermatt A & Kratschmar DV 2012 Tissue-specific modulation of mineralocorticoid receptor function by 11beta-hydroxysteroid dehydrogenases: an overview. Mol Cell Endocrinol 350 168–186.

Osorio J & Retaux S 2008 The lamprey in evolutionary studies. Dev Genes Evol 218 221–235.

Ou XM, Storring JM, Kushwaha N & Albert PR 2001 Heterodimerization of mineralocorticoid and glucocorticoid receptors at a novel negative response element of the 5-HT1A receptor gene. J Biol Chem 276 11299–14307.

Pascual-Le Tallec L & Lombes M 2005 The mineralocorticoid receptor: a journey exploring its diversity and specificity of action. Mol Endocrinol 19 2211–2221.

Pippal JB, Cheung CM, Yao YZ, Brennan FE & Fuller PJ 2011 Characterization of the zebrafish (Danio rerio) mineralocorticoid receptor. Mol Cell Endocrinol 332 58–66.

Prunet P, Sturm A & Milla S 2006 Multiple corticosteroid receptors in fish: from old ideas to new concepts. Gen Comp Endocrinol 147 17–23.

Rafestin-Oblin ME, Souque A, Bocchi B, Pinon G, Fagart J & Vandewalle A 2003 The severe form of hypertension caused by the activating S810L mutation in the mineralocorticoid receptor is cortisone related. Endocrinology 144 528–533.

Roberts BW, Didier W, Rai S, Johnson NS, Libants S, Yun SS & Close DA 2014 Regulation of a putative corticosteroid, 17,21-dihydroxypregn-4-ene,3,20-one, in sea lamprey, Petromyzon marinus. Gen Comp Endocrinol 196 17–25.

Rossier BC, Baker ME & Studer RA 2015 Epithelial sodium transport and its control by aldosterone: the story of our internal environment revisited. Physiol Rev 95 297–340.

Rupprecht R, Reul JM, van Steensel B, Spengler D, Soder M, Berning B, Holsboer F & Damm K 1993 Pharmacological and functional characterization of human mineralocorticoid and glucocorticoid receptor ligands. Eur J Pharmacol 247 145–154.

Sack JS, Kish KF, Wang C, Attar RM, Kiefer SE, An Y, Wu GY, Scheffler JE, Salvati ME, Krystek SR, Jr., et al. 2001 Crystallographic structures of the ligand-binding domains of the androgen receptor and its T877A mutant complexed with the natural agonist dihydrotestosterone. Proc Natl Acad Sci U S A 98 4904–4909.

Sakamoto T, Mori C, Minami S, Takahashi H, Abe T, Ojima D, Ogoshi M & Sakamoto H 2011 Corticosteroids stimulate the amphibious behavior in mudskipper: potential role of mineralocorticoid receptors in teleost fish. Physiol Behav 104 923–928.

Sakamoto T, Yoshiki M, Takahashi H, Yoshida M, Ogino Y, Ikeuchi T, Nakamachi T, Konno N, Matsuda K & Sakamoto H 2016 Principal function of mineralocorticoid signaling suggested by constitutive knockout of the mineralocorticoid receptor in medaka fish. Sci Rep 6 37991.

Sauka-Spengler T & Bronner-Fraser M 2008 Insights from a sea lamprey into the evolution of neural crest gene regulatory network. Biol Bull 214 303–314.

Shibata S, Rinehart J, Zhang J, Moeckel G, Castaneda-Bueno M, Stiegler AL, Boggon TJ, Gamba G & Lifton RP 2013 Mineralocorticoid receptor phosphorylation regulates ligand binding and renal response to volume depletion and hyperkalemia. Cell Metab 18 660–671.

Stolte EH, de Mazon AF, Leon-Koosterziel KM, Jesiak M, Bury NR, Sturm A, Savelkoul HF, van Kemenade BM & Flik G 2008 Corticosteroid receptors involved in stress regulation in common carp, Cyprinus carpio. J Endocrinol 198 403–417.

Sturm A, Bury N, Dengreville L, Fagart J, Flouriot G, Rafestin-Oblin ME & Prunet P 2005 11-deoxycorticosterone is a potent agonist of the rainbow trout (Oncorhynchus mykiss) mineralocorticoid receptor. Endocrinology 146 47–55.

Sugimoto A, Oka K, Sato R, Adachi S, Baker ME & Katsu Y 2016 Corticosteroid and progesterone transactivation of mineralocorticoid receptors from Amur sturgeon and tropical gar. Biochem J 473 3655–3665.

Takahashi H & Sakamoto T 2013 The role of ‘mineralocorticoids’ in teleost fish: relative importance of glucocorticoid signaling in the osmoregulation and ‘central’ actions of mineralocorticoid receptor. Gen Comp Endocrinol 181 223–228.

Tamura K, Peterson D, Peterson N, Stecher G, Nei M & Kumar S 2011 MEGA5: molecular evolutionary genetics analysis using maximum likelihood, evolutionary distance, and maximum parsimony methods. Mol Biol Evol 28 2731–2739.

Thornton JW 2001 Evolution of vertebrate steroid receptors from an ancestral estrogen receptor by ligand exploitation and serial genome expansions. Proc Natl Acad Sci U S A 98 5671–5676.

Trapp T, Rupprecht R, Castren M, Reul JM & Holsboer F 1994 Heterodimerization between mineralocorticoid and glucocorticoid receptor: a new principle of glucocorticoid action in the CNS. Neuron 13 1457–1462.

Vize PD & Smith HW 2004 A Homeric view of kidney evolution: A reprint of H.W. Smith’s classic essay with a new introduction. Evolution of the kidney. 1943. Anat Rec A Discov Mol Cell Evol Biol 277 344–354.

Wang H, Bussy U, Chung-Davidson YW & Li W 2016 Ultra-performance liquid chromatography tandem mass spectrometry for simultaneous determination of natural steroid hormones in sea lamprey (Petromyzon marinus) plasma and tissues. J Chromatogr B Analyt Technol Biomed Life Sci 1009-1010 170–178.

Williams SP & Sigler PB 1998 Atomic structure of progesterone complexed with its receptor. Nature 393 392–396.

Yang J & Young MJ 2009 The mineralocorticoid receptor and its coregulators. J Mol Endocrinol 43 53–64.

Zhang J, Simisky J, Tsai FT & Geller DS 2005 A critical role of helix 3-helix 5 interaction in steroid hormone receptor function. Proc Natl Acad Sci U S A 102 2707–2712.

